# Knowledge guided multi-level network inference

**DOI:** 10.1101/2020.02.19.953679

**Authors:** Christoph Ogris, Yue Hu, Janine Arloth, Nikola S. Müller

**Affiliations:** Institute of Computational Biology, Helmholtz Center Munich, Ingolstädter Landstr. 1 85764 Neuherberg, Germany; Department of Translational Psychiatry, Max Planck Institute of Psychiatry, 80804 Munich, Germany

**Author notes:** Corresponding authors: Christoph Ogris, Nikola Müller, Institution: Institute of Computational Biology, Address: Helmholtz Center Munich, Ingolstädter Landstr. 1 85764 Neuherberg, Germany Mail.

## Abstract

Constantly decreasing costs of high-throughput profiling on many molecular levels generate vast amounts of so-called multi-omics data. Studying one biomedical question on two or more omic levels provides deeper insights into underlying molecular processes or disease pathophysiology. For the majority of multi-omics data projects, the data analysis is performed level-wise, followed by a combined interpretation of results. Few exceptions exist, for example the pairwise integration for quantitative trait analysis. However, the full potential of integrated data analysis is not leveraged yet, presumably due to the complexity of the data and the lacking toolsets. Here we propose a versatile approach, to perform a multi-level integrated analysis: The Knowledge guIded Multi-Omics Network inference approach, KiMONo. KiMONo performs network inference using statistical modeling on top of a powerful knowledge-guided strategy exploiting prior information from biological sources. Within the resulting network, nodes represent features of all input types and edges refer to associations between them, e.g. underlying a disease. Our method infers the network by combining sparse grouped-LASSO regression with a genomic position-confined Biogrid protein-protein interaction prior. In a comprehensive evaluation, we demonstrate that our method is robust to noise and still performs on low-sample size data. Applied to the five-level data set of the publicly available Pan-cancer collection, KiMONO integrated mutation, epigenetics, transcriptomics, proteomics and clinical information, detecting cancer specific omic features. Moreover, we analysed a four-level data set from a major depressive disorder cohort, including genetic, epigenetic, transcriptional and clinical data. Here we demonstrated KiMONo’s analytical power to identify expression quantitative trait methylation sites and loci and show it’s advantage to state-of-the-art methods. Our results show the general applicability to the full spectrum multi-omics data and demonstrating that KiMONo is a powerful approach towards leveraging the full potential of data sets. The method is freely available as an R package (https://github.com/cellmapslab/kimono).

## Introduction

Over the past decade, high throughput techniques enabled the possibility to study biological mechanisms on a large scale and in a cost-efficient manner. The resulting tremendous increase of omic data has the potential to provide deep insights into complex biological processes being orchestrated by the interplay of a diverse network of biomolecules. The number of bi- or multi-omics data sets increase, yet combined analysis is not yet well explored. Several reviews (Pinu et al. 2019; Hasin, Seldin, and Lusis 2017; Huang, Chaudhary, and Garmire 2017) discuss the potentials of multi-omics data algorithms directly pointing out the methodological gap of identifying and analyzing cross omic relations. One exception is the field quantitative trait (QT) analysis (Heinig et al. 2017; Zhernakova et al. 2017). Here two types of information are linked, for instance, the genetic or methylation site with gene expression, in order to explain variation in complex traits. However, integrating multiple levels simultaneously is still an ongoing challenge.

The most common integration approach so far is to explore available omic levels independently and search for common significant features afterwards (Sinkala, Mulder, and Martin 2020). This level-by-level analysis not only misses features with low signal but also ignores the complex ‘cross-omic’ interplay and might subsequently cause misinterpretation of the data (Domenico et al 2015, Schmitt et al 2013).

Recently, sophisticated latent factor-based omic integration approaches have been introduced (Ronen, Hayat, and Akalin 2019; Argelaguet et al., n.d.). These methods infer lower-dimensional representations, latent factors, of the original high dimensional multi-omic data space. Even though these can represent certain patterns of the data, it is still unclear how many latent factors are needed to completely describe the complex data structures (Rares-Darius Buhai et al. 2019). There seem to be many advantages of using these methods for reducing dimensionality. However, it is often difficult to infer the biological meaning from latent factors (Argelaguet et al., n.d.). Moreover, it is not possible to identify inter- and intra-associations between latent factors or the features within different omic levels.

These issues are accounted for by network inference approaches, reconstructing the interactome in the form of a network describing the interplay of all features within the data. The assembled network structure can then be used to identify key features, like modules and pathways, underlying a disease. The most common straightforward network inference approach is to use pairwise correlations, linking all features which are significantly correlated. These networks are often hard to interpret since they tend to be too densely connected (Krumsiek et al. 2011). This can be accounted for by using machine learning approaches like graphical random forests (Lee and Hastie 2015; Zierer et al. 2016). But, these methods are in need of large amounts of samples. To overcome this, we developed miRlastic (Sass et al. 2015) which facilitates prior knowledge to increase the performance for high dimensional and low sample size data analysis. MiRlastic creates a regression model for each mRNA with, previously as target reported, miRNA species. Next, aggregating miRNA and mRNA species from all regression models assembling an mRNA-miRNA interaction network. Based on this network, we were able to identify, yet undiscovered, functional miRNA target clusters.

Our novel multi-omic network inference approach, named KiMONo, vastly expands our miRNA-mRNA integration idea (Sass et al. 2015) now allowing a simultaneous integration of multiple omics levels, for example, variants, methylation, gene expression, proteomics, biological information. KiMONo uses a regularized regression model for each gene and leverages various prior information to reduce the high dimensional input space. KiMONo uses, per default, a sparse group lasso as a penalty model. This allows for a bi-level selection, penalizing each information level as a group but also penalizing within each level. Aggregating these models, all non-zero features are linked to their gene and overall assemble the heterogeneous multi-level network., hence it does not restrict K to any specific omic types.

## Materials & Methods

### Network inference with KiMONo

Our novel method KiMONo infers heterogeneous/multi-level networks to identify key features and better understand the interplay of various omic levels in a biomedical setting where matched multi-level omics data is available. For efficient and biologically reasonable inference, KiMONo makes use of prior knowledge to link available data to the transcriptomic level, thereby generating a network representation of prior knowledge (Figure 1). Within this prior network, each node represents a feature of the multi-omic input data, connected to a gene. Links between nodes indicate prior-defined associations. Such an association can range from an experimentally validated genetic or predicted gene-protein interaction up to simple annotations between genes coding for proteins. On top of using only these first-order links, KiMONo can also process second-order links as prior. These can be generated by interconnecting the transcriptome and linking features of associated neighbours to a gene.

**Figure 1:**
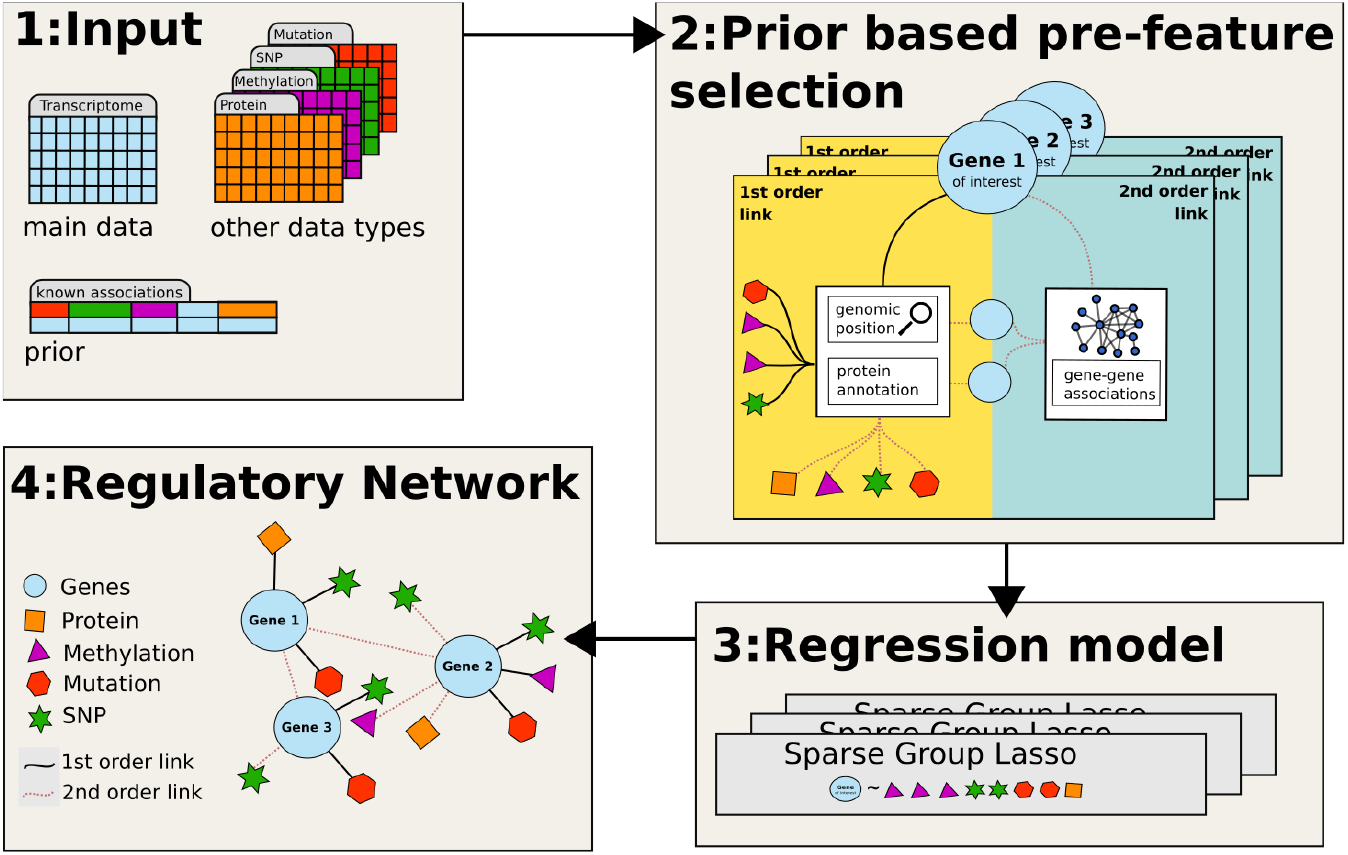
Workflow: First, KiMONo can integrate measurements of any number of different data types. Samples need to be matched, indicated by the same number of columns of the matrices. The method also requires a prior in form of binarized mappings between the data types. Second, based on these mapping the algorithm creates an input matrix *X* for each gene loading required measurements that defines the gene-based initial feature space. Third, by using the genes expression as *y*, the algorithm further runs a regression model based on Sparse Group LASSO penalization. Fourth, all gene-based models are merged to compile a multi-level network containing features from all input sources as nodes and links for all non-negative regression coefficients between them define the data specific relation.

Once the prior network is established, KiMONo optimizes an individual penalized regression model for each node within the transcriptomic level. Here the gene expression represents the criterion variable *y* while the input matrix *X* is assembled by the standardized feature vectors of connected nodes within the prior network. KiMONo uses the sparse group LASSO (Simon et al. 2013) regression approach which penalizes within and between predefined groups of features. By performing this ‘bi-level’ selection, we can account for different underlying distributions between the features which originate from using multiple data types. Within sparse group LASSO, the parameters *α* denotes the intergroup penalization while *τ* defines the group-wise penalization. KiMONo approximate an optimal parameter setting via using the Frobenius norm (Sass et al. 2015). To be more specific, *α* is approximated by the mean Frobenius norm of all groups while *τ* is estimated by the frobenius norm within each group. The global LASSO parameter *λ* was estimated via 5-fold cross-validation, using the mean squared error as loss function.

KiMONo further uses the fitted models, of all nodes within the priors transcriptome level, to assemble a multi-level omic network. Within this network, nodes represent features of the input data, like genes, and connections between them are defined by the optimized *β* coefficient. Furthermore we assign each gene node a confidence score by its modeled *R*^2^.

### The Cancer Genome Atlas data and prior

The cancer cohort data was obtained via one of the most comprehensive multi-omic data sources, The Cancer Genome Atlas (*TCGA*) data portal (Weinstein et al. 2013). We here focused on *TCGA*’s well described *PanCancer* data. This collection contains multi-omic data sets of 4926 samples describing 12 different cancer types - *acute myeloid leukemia (191* samples), *bladder urothelial carcinoma (135* samples*), breast invasive carcinoma (871* samples*), colon adenocarcinoma (421* samples*), glioblastoma multiforme (580* samples*), head & neck squamous cell carcinoma (309* samples*), kidney clear cell carcinoma (496* samples*), lung adenocarcinoma, lung squamous cell carcinoma (344* samples*), ovarian serous cystadenocarcinoma (563* samples*), rectum adenocarcinoma (164* samples) and *uterine corpus endometrioid carcinoma (495* samples). The portal provides open access to highly preprocessed ‘*level 3’* data of five omic characterizations, *Proteome (~130 proteins)*, *Transcriptome (~ 16115 transcripts)*, *Copy Number Variation* (~*84 CNV*), *Mutation (~39675 positions)*, *Methylation (~2043 sites)* but also phenotype information in the form of *Clinical* data (*4 variables*). In our analysis we only included samples which were measured across all 5 omic levels, restricting the data sets to 2036 patients across 11 cancer types (see Sup Fig 1). Beside binarizeing the *Clinical* feature ‘sex’ we also standardized all input features.

In order to assemble the prior knowledge networks for the *panCancer* cohort, we used both first- and second-order links to connect the *Transcriptome* to all information levels. First-order links to the *Proteome* were generated via the *bioMart* annotation resource. First-order links to *CNV* and *Methylation* were generated via a genomic position-confined prior. Here we used Bioconductor’s R packages *Homo.sapiens*, *GenomicRanges (Lawrence et al. 2013)* and *FDb.InfiniumMethylation.hg19 (*Triche et al 2014*) to link* copy numbers and methylation sites within a 500kb range to genes of interest. The *Mutation* data type was already projected to gene identifiers, hence there were no additional preprocessing steps needed. Furthermore, we used experimentally validated associations from *BioGrid* (Oughtred et al. 2019) to create links within the *Transcriptome*. Additionally, we added second-order links to increase the coverage of individual gene models. This was done by connecting genes to first-order linked features of gene neighbours. In a final step, we also connected all features within the *Transcriptome* to the *Clinical* features.

### Major Depressive Disorder data and prior

The Major Depressive Disorder (*MDD*) cohort consists of 289 caucasian individuals, 160 healthy controls and 129 patients diagnosed with major depressive disorder. Recruitment strategies and further characterization of the *MDD* cohort have been described previously in (Arloth, Bogdan, et al. 2015) and (Zannas et al. 2015). Three levels of omic information, comprising of the transcriptome, methylome and genotype, as well as biological information, were measured for 107 out of 289 individuals, consisting of 33 females and 74 males, distributed over 64 controls and 43 patients. Details on the omic preprocessing can be found in (Arloth, Bogdan, et al. 2015) and (Zannas et al. 2015).

For generating the prior knowledge first-order links, we annotated gene expression probes and gene symbols using the Re-Annotator pipeline (Arloth, Bader, et al. 2015) based on GRCh37 (hg19) RefSeq. Additionally, we annotated methylome, the CpG site probe, and the transcriptomes gene symbol to sequences positions by performing a re-alignment using Bismark (Felix Krueger 2011). Furthermore, we connected the genes to SNPs and methylation sites within a distance of 10 kbp and 500 kbp, respectively. Second-order links were created between genes via a ‘guilt-by-association’ approach using the BioGrid database. Furthermore, we connected genes with their associated genes methylation site generating, introducing second-order linked methylation sites.

### Performance test

To assess goodness of fits on every gene-level model, we use the r-squared metric measuring how much of the variance of the expression can be explained by the model. We calculate for each model the explained sum of squares *ESS*, defined as 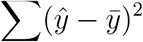, and total sum of squares 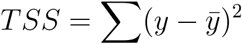. Here, *y* represents the true and 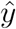 the predicted gene expression. The amount of variance explained is then given by *R*^2^ = *ESS/TSS*.

In order to approximate each information level contribution to the *R*^2^, we dissect the *R*^2^ and calculate a 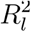 for each information level. This is done by calculating the 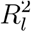 via 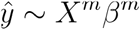 and *y*_*new*_ = *y* − *X*^*n*^*β*^*n*^. Here *m* defines all features within level *l* and *n* denotes all other features. *y*_*new*_ and 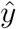 were further used to estimate a 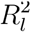 which approximates the sole performance of *l*. Finally, 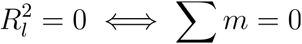, which sets the performance of levels without selected features to 0.

### Prior expansion by 2nd order links

Overall, we compared two different prior strategies. On the one hand, a prior solely based on the genomic location and annotation databases. Here we annotated protein, methylation, mutation and clinical information to the transcriptome level. On the other hand we generated a prior also including second-order links using the BioGrid(Oughtred et al. 2019) resource. We not only interconnected the transcriptome but also all other layers.

We used the *panCancer breast invasive carcinoma* network models as test scenarios, investigating the impact of the different prior strategies. To evaluate how well each strategy performed, we compared the performance of the models which explained at least 10% of the variance within the data and the coverage of the inferred networks.

### Robustness to measurement noise & low sample sizes

To benchmark the performance on small data, we simulate data sets with shrinking sample size. Therefore we used the *TCGA breast invasive carcinoma* data and randomly reduced the amount of samples. We repeated each simulation 20 times (except for the 100%). The final test cases included 10%, 30%, 50%, 70% and 100% of the data. For each generated dataset, we excluded features with *σ* = 0.

We followed a similar strategy for benchmarking the robustness of the method with respect to noise. Here we simulated test sets by decreasing the signal to noise ratio. All simulated sets were created using a subset of the *panCancer breast invasive carcinoma* data. Random noise was generated using Gaussian noise, 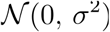 with increasing *σ*^2^. Here we simulated noise with *σ* ∈ {0,0.2,0.4,0.6,0.8,1}.

For both, we used the above described *R*^2^ and 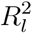 metric to evaluate the models’ performances, excluding all models *R*^2^ < 0.1.

### Quantitative trait analysis

We compared the quantitative trait analysis results of KiMONo to the state-of-the-art pairwise analysis tool, matrixEQTL. Here we used both methods to detect expression quantitative trait loci (eQTL) and expression quantitative trait methylation sites (eQTM) genes within the MDD data set. For the matrixEQTL calculation, we focused on cis-eQTL and cis-eQTM windows of 10 kbp and 500 kbp distance, respectively. Further, we corrected the expressed genes for the covariates, BMI, age, sex and status of the diagnosis, with significance threshold set to *FDR* < 0.05.

In the case of KiMONo, eQTL and eQTM genes are identified via the inferred cross-layer interactions between genes and methylation sites and SNP’s. Here, robustly inferred results were defined as models with *R*^2^ ≥ 0.1 and the respective cross-layer association of |*β*| ≥ 0.2.

### Network statistics

We transformed the directed links to undirected associations, generalizing the multi-layer directed network to a simpler single-layer association network representation. To show that the generalized network structure is, like most biological networks, scale-free, we used goodness of fit test to evaluate if the node degree follows a gamma distribution (X. Wang, Gulbahce, and Yu 2011). Furthermore, we used the *betweenness centrality* to estimate the importance of nodes within the single-layer network. The *betweenness centrality* is defined as 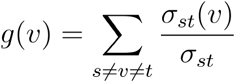. Here *σ*_*st*_(*υ*) defines the shortest path between node *s* to node *t*, passing node *υ*.

### Data access

The *panCancer* data is publicly available via the *TCGA* data portal (downloaded May, 2017). A list of the sampleIDs and cancer types which contained all 5 omic levels can be found in Sup Data 1. The transcriptomic and epigenomic information layer of the MDD cohort can be found at GEO GSE64930 and GSE74414, while the snp data cannot be provided due to patient privacy regulations.

### Used software

KiMONo is freely available via the R package https://github.com/cellmapslab/KiMONo. We used the Bioconductor’s R packages *Homo.sapiens*, *GenomicRanges* and *FDb.InfiniumMethylation.hg19* to generate the annotation between various omic types within the TCGA data. Furthermore we used Re-Annotator pipeline (Arloth, Bader, et al. 2015) based on GRCh37 (hg19) RefSeq and Bismark (Felix Krueger 2011) to annotate the MDD data. For state-of-the art eQTL analysis we applied matrixEQTL (version 2.3). Pathway annotations were performed via pathwaX (Ogris, Helleday, and Sonnhammer 2016).

## Results

With the versatile tool KiMONo at hand, we seek to investigate its capabilities and limitations first on one of the largest and most comprehensive multi-omics dataset collections available today. The panCancer dataset of *TCGA* was downloaded and processed before data of every cancer type was subjected to KiMONo network inference with default parameter. Subsequently we have been using the *panCancer breast invasive carcinoma* dataset for in depth analysis, as it is one of The subset of TCGA’s *breast invasive carcinoma* data contains 604 patients with in sum 57 966 features measured across 5 data types. Secondly, we used multi-omic dataset for a common complex disorder, MDD. It consists of 289 individuals with in sum X features measured across 4 data types in peripheral blood cells.

### Improved performance using second-order links

First, we asked whether combining first- with second-order links increase performance of gene-based models. We also evaluated the performance using all features at once without any prior knowledge, however, the algorithms inferred only empty networks due to the high dimensionality of the data.

Evaluation of KiMONo, using only first-order prior resulted in 5349 inferred gene models with 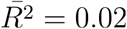. Only 96 gene models were evaluated with an *R*^2^ ≥ 0.1 and had an average 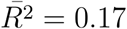. Adding second-order associated features, we were able to learn models for 9480 genes having an average performance of 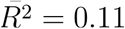 and 3150 gene models were created *R*^2^ ≥ 0.1 with and 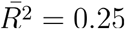. Furthermore, we had a closer look at all models with *R*^2^ ≥ 0.1 and evaluated the impact of the amount of different information layers within a model (see Figure 2.A). One can observe that a diverse set of data types increases the model performance. Moreover, we can see that there is no model relying on a single information layer alone. Models based on features of two information layers show an average 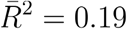 while five information layers increase the variation explained to 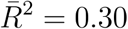. This is also directly mirrored by the number of features used, see Figure 2.B. Here the majority of two-leveled models are based on an average number of 3.6 related features while models using five layers detected on average 77.5 associations.

**Figure 2:**
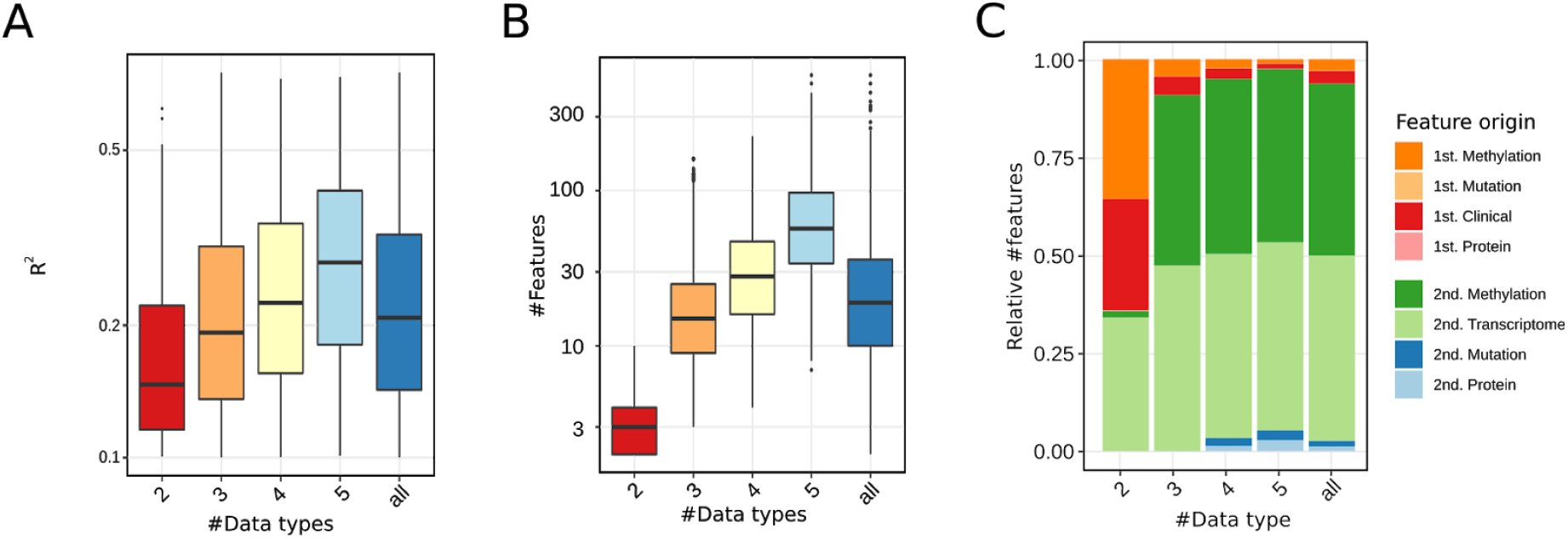
A) Boxplots of performances of models relying on 2, 3, 4, and 5 different data types/omic levels. The last boxplot denotes the overall performance. B) Number of features used if a model includes different data types. Here the last box refers to the general number of features used. C) Barplots visualizing the amount of omic types included in models. Here orange-red refers to 1st order linked features while green and blue visualize 2nd order features.

### Performance on small sample-sized data

One of the biggest challenges for data analysis methods is to be applied to data that has a lot of features and only a small amount of samples. This is, in particular, relevant for multi-omic data sets which often comes with thousands of features but just a few samples. Our data simulations of small sample sizes resulted in 140 test sets, based on the *breast invasive carcinoma* data provided by *TCGA*. Each set is a representative subset of samples of the original set. Using KiMONo we assembled a network for each of these test cases (see Figure 3.A). Applying KiMONo on all *breast invasive carcinoma* data samples resulted in an inferred network coverage of 7223 gene models (of 11530) with *R*^2^ > 0.01 (see Sup. Figure 3.A). While reducing the sample size to 30 we were able to retain 5338 of the initial genes, losing only 1885 models. In the case of higher-performing models (*R*^2^ > 0.1) we were able to find 80% of the initial models for 5% of the initial samples.

**Figure 3:**
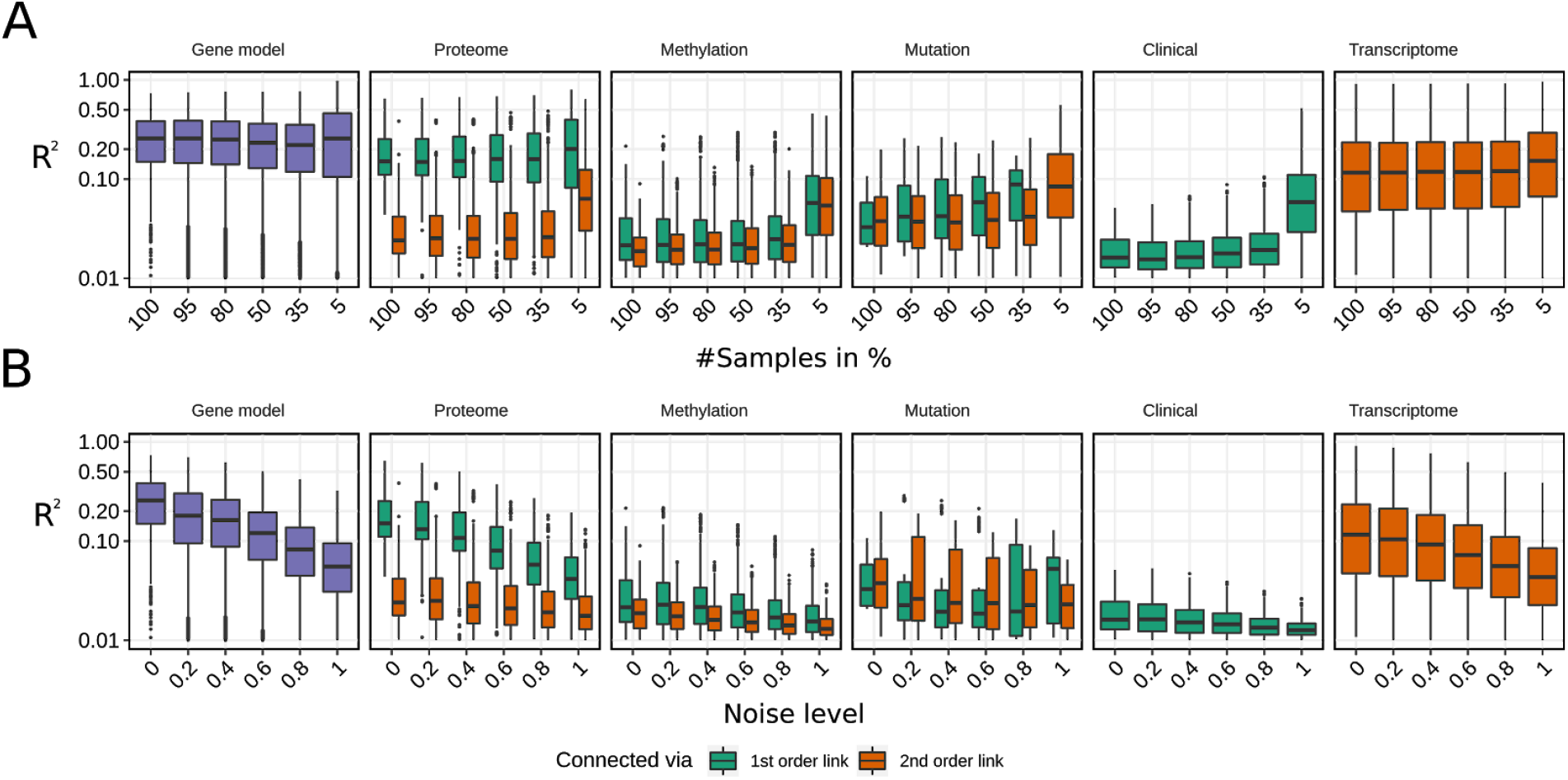
Robustness benchmark for A) different sample sizes and B) noise levels. First set of box plots (purple) shows the performance, *R*^2^ (log scaled x axis) of inferred gene models using all available information layers. All following sets describe the stand-alone performances of Proteome, Methylation, Mutation, Clinical and Transcriptomic information layers. 1st order links (green) and 2nd order links (orange) are benchmarked separately. Note, Clinical and Transcriptome information consists of only 1st and 2nd order links. A) Data sets with different sample sizes were generated using 10% - 100% of the 604 breast invasive carcinoma samples. B) Different test data sets were simulated by adding Gaussian noise with increasing variance. Here, the noise level reflects the σ for six intensities.

To evaluate the overall performance we excluded genes models which explained less than 1% of the gene expression variance (*R*^2^ < 0.01) and restricted the benchmark set further to 932 genes which have been also present in the 30 sample size test cases (an unrestricted view can be found in Sup. Figure 3). Removing samples also decreased the variance of many features which indirectly decreased the overall dimensionality. Using 30 (5%), of the 604 samples, reduced the number of features from 57 966 to 15 632 features.

Comparing the overall results we can show that KiMONos performance is stable for small as well as large sample size data. This reduction of complexity is reflected by a slight increase in performance between the 35% and 5% test cases, from 0.24 to 0.29.

Following our approach of dissecting the overall *R*^2^, see Materials and Methods, we were able to estimate the importance of the individual levels as well. The most informative sole information layer is the *Protein* information with and average *R*^2^ of 0.23 followed by the second-order linked *Transcriptome* information with an average *R*^2^ of 0.18. The sparse *Mutation* data seems to improve its performance with smaller sample sizes whereas *Clinical*, *Methylation* and second-order linked *Protein* information seems to contribute the least. When comparing the results between the different sample sizes, one can observe that the mutation layer constantly improves the performance by an *R*^2^ of 0.7 for reduced sample sizes with lower dimensionality while all other information layers slightly decrease in performance.

### Performance on noisy data sets

To simulate noisy data set we used the *breast invasive carcinoma* data set as a blueprint. Following the simulation approach described in Materials and Methods, we generated 100 different data sets across five noise levels. Using KiMONo we inferred networks for all test cases. In terms of network coverage, we can see a more reduced coverage in noisy data than compared to the reduced sample sizes (see Sup Figure 2.B). KiMONo is able to retrieve more than half (4071) of the initial models with *R*^2^ > 0.01. Looking at higher-performing models (*R*^2^ > 0.1) we can see that the gene coverage of 3147 models drops to 463.

This performance drop can also be observed by evaluating the overall gene model performance. Here we only evaluated models explaining at least 1% of the variance within the gene expression. The most drastic impact of noise can be observed at the 1st order linked *Proteome* and 2nd order *Transcriptome* data. In the *Proteome* data, the performance drops from 0.21 to 0.05 while the *Transcriptome* decreases from 0.17 to 0.6.

The overall average *R*^2^ = 0.28 is decreased to 0.05. after adding Gaussian noise with *σ* = 1. Similar to the previous performance test, we can again observe that all other information layers have similar overall trend. Information layers that already start with a relatively low *R*^2^ like *Methylation (0.03)* and *Clinical* (0.02) layer. Here the performance was reduced to 0.02 for *Methylation* and 0.01 for *Clinical* data respectively.

### Multi-layer pancancer networks

To exemplify the data analysis power of KiMONo on multi-layer data we inferred networks for 11 cancer types. As a post-processing step, we excluded all models for which *R*^2^ < 0.1 and also excluded links within the network with a weight smaller than |*β*| < 0.02.

The final networks have on average 26.343 links and 3158.2 nodes (see Figure 3.A). Test of fit for the degree distribution being gamma distribution resulted in a *p* < 2.2*e* − 16. For each network, we ranked the nodes based on the node betweenness of centrality and selected the top 100 features. Comparing these sets shows that 88% of the top 100 features are occurring in at least two of the cancer types. All genes which are identified as important across all 11 cancer networks have been previously linked to cancer by several studies (see Sup. Table 1). We further used all those genes for pathway annotation using the tool pathwaX (Ogris, Helleday, and Sonnhammer 2016). Here the top enriched KEGG pathway is the cancer-related *Chronic myeloid leukemia* (FDR=1.45e-37) pathway followed by *Pathways in cancer* (FDR=6.3e-35). Furthermore, we were also able to identify 345 features which were uniquely identified to each cancer type. For instance, the methylation site cg00103783 (chr17:7.583.931), mapping to *MPDU1* gene, was only detected as important within the *head & neck squamous cell carcinoma* network. Here (Ceder et al. 2012) introduced MPDU1 as a potential biomarker for HNSC. Within the breast invasive carcinoma network, all three genes are among the top 20 features, lead by *age* and *UBC* which has been identified as an oncogene by (Ohta and Fukuda 2004) (see Figure 4). Using these top 20 genes for pathway annotation gives a clear picture of cancer-related KEGG pathways, i.e.: KEGG *Pathways in Cancer* (FDR = 2.94e-44) is the top enriched pathway, followed by *Hepatitis B* (FDR=2.51e-39) and *Cell cycle* (FDR=2.3e-38). Both, *Cell cycle* and *Hepatitis B*, are known breast cancer-related pathways (Yeo et al. 2003), (Catzavelos et al. 1997). However the *Breast Cancer-specific* KEGG pathway ranks on place 14 (FDR = 5.0395e-33) among all enriched pathways. Another interesting result is the inferred *Glioblastoma multiforme* (GBM) network. Even though *GBM* is one of the rarest cancer types it is also one of the most lethal ones having a survival time of 14-15 months after diagnosis (Hanif et al. 2017). The *GBM* data set is relatively small including only paired data for 61 patients with 58051 features across 5 omic layers. Nevertheless KiMONo inferred 112945 links between 9341 nodes. Even though the top 20 features are not as densely connected as in the previous example we were able to link *CTNNB1, HIF1A*, *HDAC1 and EWSR1* to *survival time.* Beside *EWSR1* all have been reported as survival time related in GBM ((McCord et al. 2017), (Liu et al. 2015). Interestingly (Bridge et al. 2019) show that in GBM, EWSR1 is often fused with *PATZ1* which was in the past related to worse survival rates (McCord et al. 2017; Guadagno et al. 2017).

**Figure 4:**
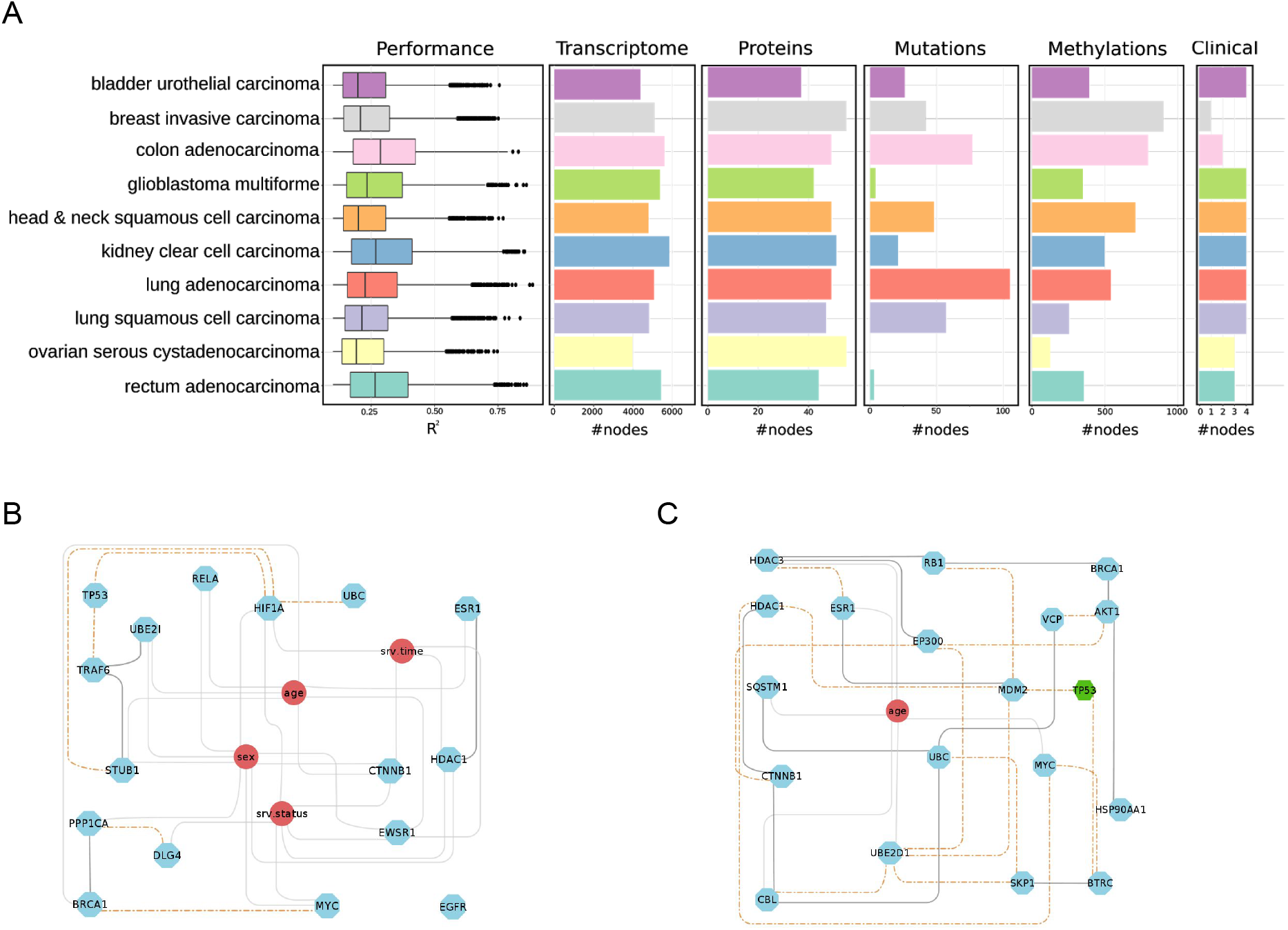
A) Performance on all gene models (*R*^2^ > 0.1) inferred by KiMONo followed by amount of gene models and amount of features selected in the proteomics, mutation, epigenetic and clinical data layer. B) Subnetwork of top 20 features (highest betweenness of centrality) within the inferred breast invasive carcinoma network 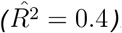. Within the network, we can find nodes originating from mRNA (blue), mutation (green) and clinical (red) feature space. The edges denote first order edges (grey), first and second-order combined (black) and las only second-order connections (dashed orange). C) Subnetwork of the top 20 features in Glioblastoma Multiforme 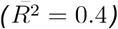.

### Multi-layer MDD network

Even though the TCGA is one of the most comprehensive multi-omic datasets available, we wanted to evaluate our method on a more complex type of disease, like MDD. While progress has been made in understanding the pathomechanisms involved in MDD, success in translating findings into clinical practice has been limited (Kapur, Phillips, and Insel 2012). To this end, studies have been largely focused on single-level omics, like GWAS (Howard et al. 2019)) and multi-level omics are relatively new (Anderson et al. 2020; Arloth, Bogdan, et al. 2015). Therefore, making successful inference of a multi-omic cross-talk regulatory network is of importance to better understand the depression phenotype.

For this purpose, we applied KiMONo on a patient cohort, consisting of 107 healthy individuals and patients. There were 4,247,909 imputed SNPs, 12,418 transcripts and 320,481 methylation sites available for the evaluation of our method, after filtering for the 25 % of methylation sites with the least variance. Biological information such as BMI, age, sex and status of the diagnosis and cell type composition were always taken into account for network inference.

After filtering out *β* coefficients between −0.02 and 0.02 as well *R*^2^ ≤ 0.1 as values, the final MDD network, comprised of 5,791 gene models whose gene expression was explained up to over 0.8 *R*^2^ with the median at 0.256. As predictors, we uncovered 6,894 methylation sites and 3,404 SNPs as first-order links, as well as 5,041 gene transcripts and 4,201 methylation sites as second-order links. In addition, all of the biological covariates were found across the whole network. (Figure 5.A and B)

**Figure 5:**
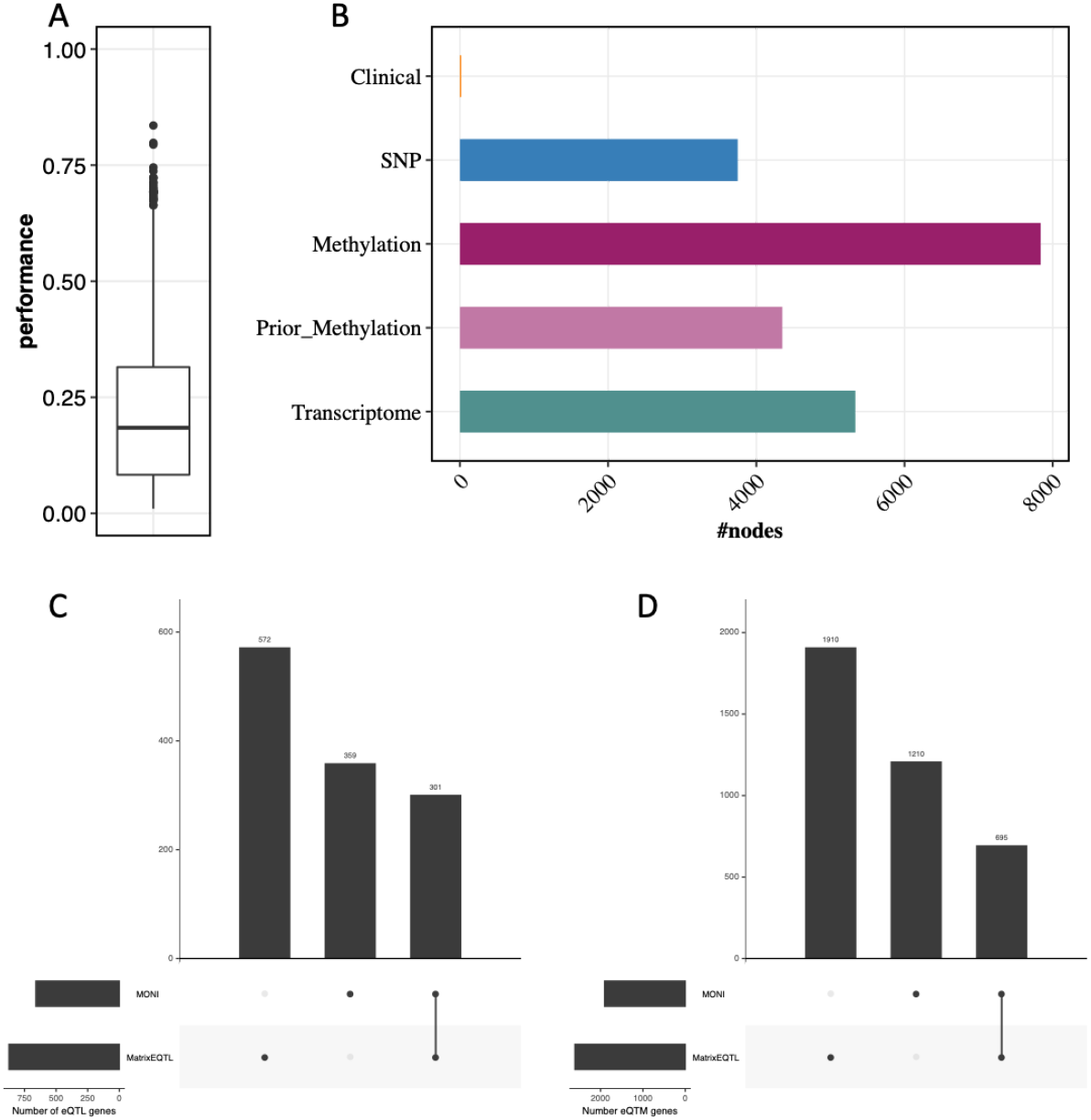
A) MDD network Performance on all gene models n=5,791 inferred by KiMONo after filtering for −0.02 < *β* < 0.02 coefficient and *R*^2^ > 0.1 B) Composition of retained features deriving from omic levels of first-methylation and SNPs, as well as and second-order methylation, SNPs, gene expression and biological clinical features; Comparison of C) number of eQTL genes and D) eQTMs gene derived from KiMONo and matrixEQTL.

To compare with state-of-the-art methods, we identified eQTL and eQTM genes using pairwise models and set them into context to the findings of KiMONo. Using the same proximity restrictions for the MatrixEQTL and KiMONo, we found 873 and 660 eQTL-genes, respectively, overlapping in 301. Further, we found an overlap of 695 eQTM genes, with 1210, more than double found with KiMONo. Nearly all genes found in the overlap or only by KiMONo were further explained multivariate models by information from other omic-layers of methylations, SNPs and gene expression.

The top 20 genes were identified with the highest betweenness measure with the performance found to be higher than the average models across the whole data set. *R*^2^ ranged from 0.202 to 0.798 with a median of over 0.491, while the average across all models was 0.256 (Figure 6.A). Further, features selected by the penalty model represented information from many different omic-information levels, across methylation, SNP, gene expression as well as biological clinical information. Methylation sites possessing long-distance effects, gene expression associated over indirect links, and biological data were consistently present for the top 20 hits (Figure 6B).

**Figure 6:**
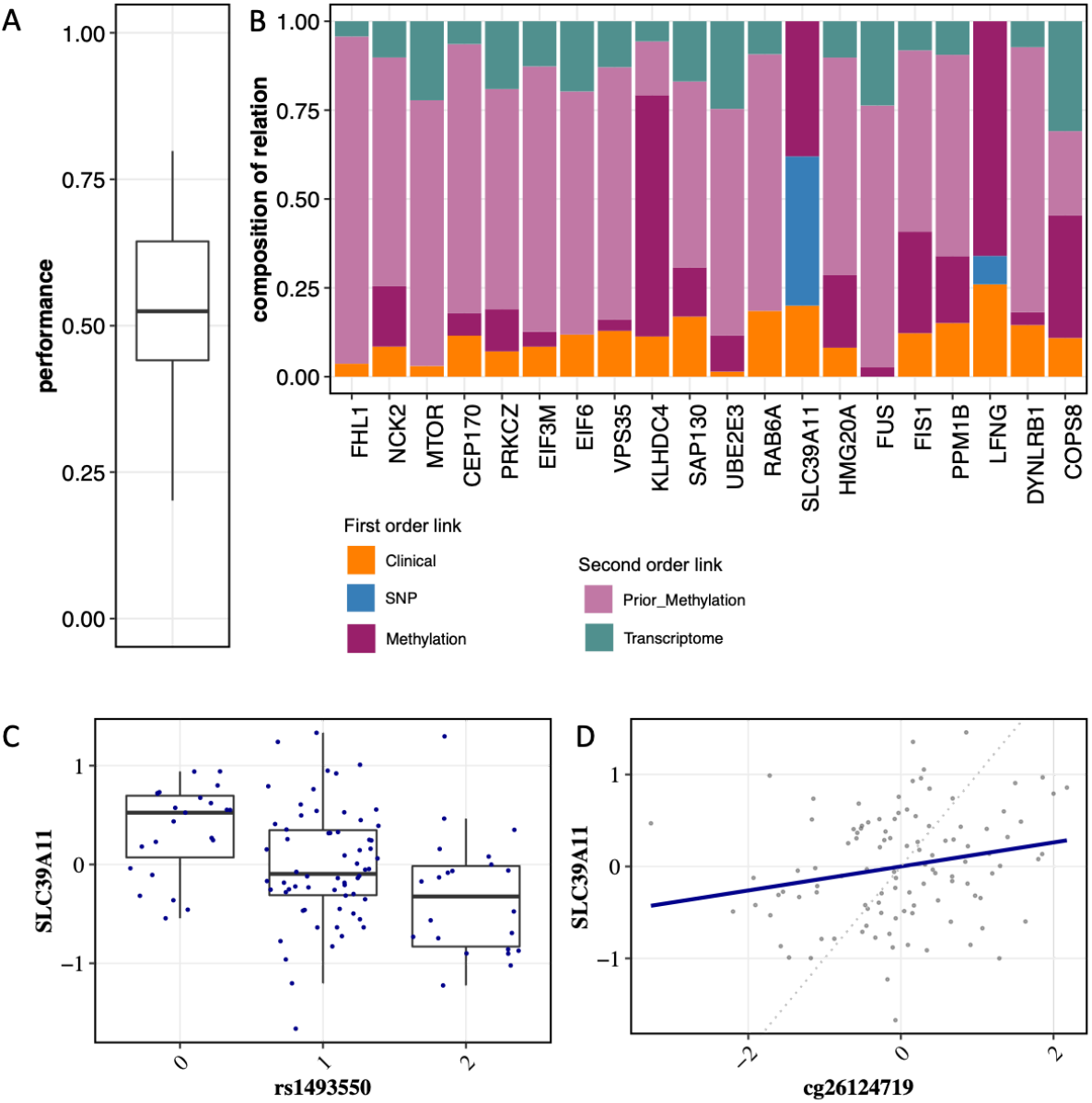
A) Performance of n=20 genes with the highest betweenness and B) its composition of retained features deriving from omic levels for each gene. Gene expression with possible influence by C) SNP and D) methylation site found with KiMONo but not with MatrixEQTL; the dotted line represents a correlation of 1.

The potential of our method becomes apparent while looking at connections found through KiMONo but not pairwise models of MatrixEQTL. After correcting for residual effects of all other features in multilevel models, the connection between the expression of SLC39A11 (Solute Carrier Family 39 Member 11, chromosome 17) and SNP rs1493550 and methylation site cg26124719 located both in an intron became clearly resolved (Figure 6.C and D).

Half of the top 10 hits have been previously linked to depression or pathways involved in the pathogenesis of the disease (see Table 1). Here the top enriched KEGG pathway is endocytosis (FDR=4.832e-8) which play a major role in synaptic plasticity, which is an important component in disease development of stress-related disorders, like MDD (Hua et al. 2013), (Duman et al. 2016). The second important pathway is autophagy (FDR=2.606e-6) an essential pathway for the central nervous system and studies have shown the effects of antidepressant treatments on autophagy (Gassen and Rein 2019). Interestingly, among the top 10 pathways is Axon guidance (FDR=1.054e-3), which has been shown to be a strong risk factor for depression, as stress may affect brain structure and function ((Breen et al. 2018), (Engle 2010).

**Table 1:**
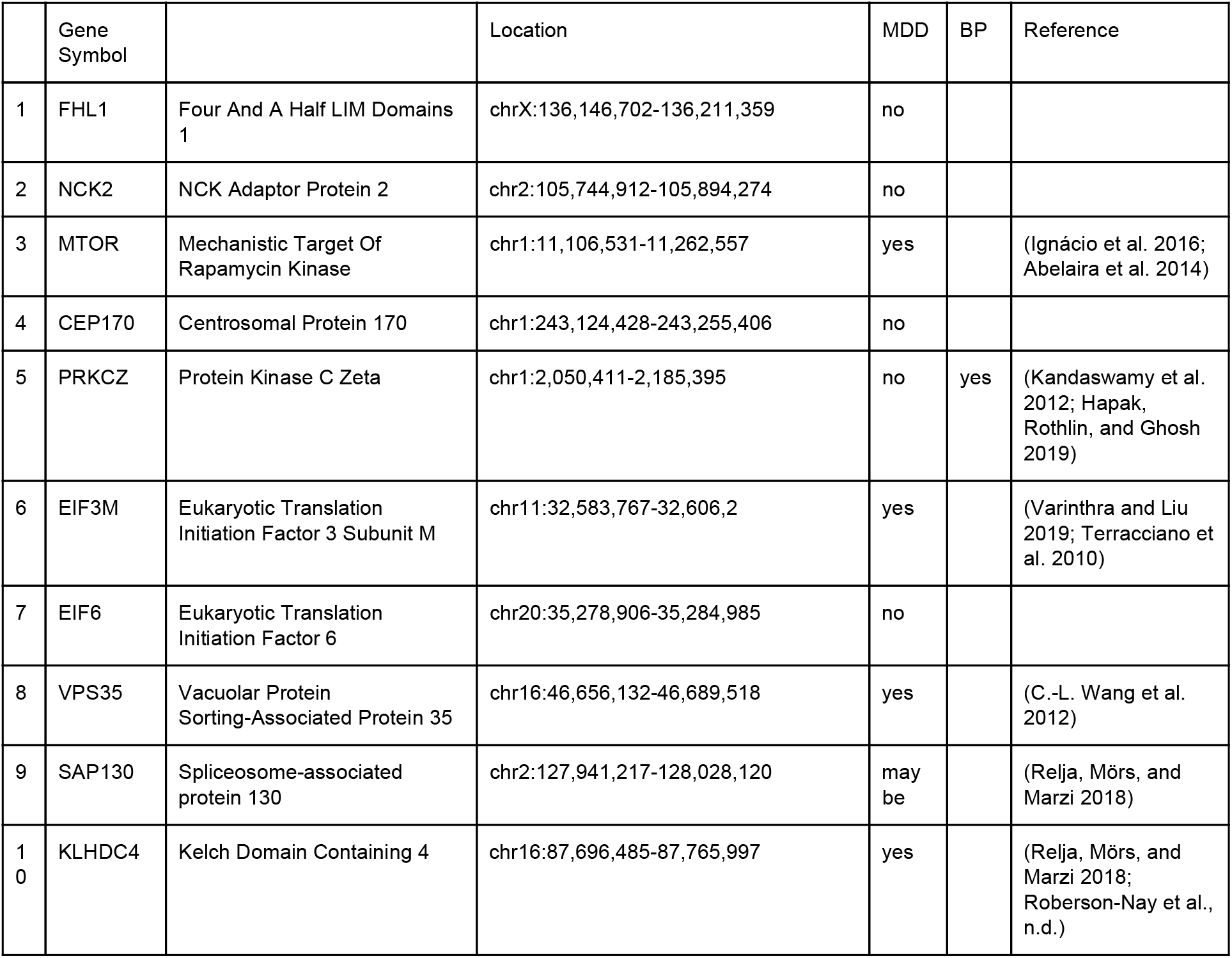
Top 10 genes of most important nodes within Major Depressive Disorder (MDD) data. Ranking was derived via the nodes betweenness of centrality.

## Discussion & Conclusion

We presented KiMONo –- a novel prior Knowledge guided Multi-Omics Network inference method. By leveraging prior knowledge, the algorithm builds a statistical model for each gene, selects the most predictive features and uses these to assemble a heterogeneous network. Within this network nodes represent features of the input sources and links define disease- or context-specific relation between them. KiMONo was specifically designed to work on low sample size sets with high-dimensional data originating from a variety of information sources.

We used TCGA data, one of the biggest collections of multi-omic data, as the main evaluation set. However, the publicly accessible data was lacking quality and information depth. For instance, mutation and methylation data were only available in a sparse binarized form. We reasoned that KiMONo enhances the signal by combining various data sources and is therefore well suited for the analysis for this data format. Nevertheless, we also performed our tests on less preprocessed data describing MDD. Even though this data set has a higher dimension, we were also able to reproduce the performance insights we gained from the TCGA data (see Sup Figure 5).

In our robustness tests, we showed that, reducing the number of samples barely affected the overall performance of KiMONo. When investigating the performance contribution of the mutation features alone, there was even a slight performance increase for low sample sizes. Even though it might be the sole effect of overfitting, we showed that it only occured for sparse binarized data. Hence, removing samples from this sparse matrix directly resulted in setting some features *σ* = 0. Therefore, we not only removed samples but also shrank the feature space. which in turn resulted in less predictive models having slightly better regression performances.

In contrast, we found that the method was more sensitive to noise in the data than performing on small sample sized data sets. When increasing the simulated noise, it resulted in a rapid decrease in correctly predicting the gene expression level, as opposed to a moderate decrease when reducing the amount of samples. This finding highlights the importance of high-quality sequencing of omic data for robust inference of regulatory networks.

Next, we showed that KiMONo was able to find many of the eQTL and eQTM genes (34.5 and 26.7%) that were uncovered by MatrixEQTL using pair and level-wise tests. In addition we found further associations, complementing MatrixEQTL, when deriving regulatory networks in context with all features from all omic levels. It is possible that these features can only be detected when taken into account the context of the underlying omic-crosstalk. Across all top hits in the MDD dataset (Table 1), we observed that relationships from 2nd-order linked genes and methylation sites play an important role. For example, gene SLC39A11 beeing identified as eQTL and eQTM gene to SNP rs1493550 and methylation site cg26124719. Our results indicate that KiMONO is a powerful method to discover these long-distance and indirect relationships while establishing regulatory networks.

In addition to incorporating second-order links, we also showed the advantage of multivariate models derived from various omic-layers by uncovering relationships that were not found in pairwise models. After correcting for residual effects of every feature except for the one of interest, the connection became clear (Figure 6C and D). Our approach allows the uncovering of many more effectors by accounting not only for the covariates but also all other features in a complex multi-omic context.

Applying KiMONo on both TCGA cancer types and MDD, we were able to find previously reported genes that matched well with the underlying disease setting (see Figure 6.B/Table 1). This provided a good evaluation of our method. Among the top hits we also identified genes that have not yet been reported in relation to the studied phenotypes. These genes could be essential for further exploration of the disease mechanisms for better understanding of the underlying molecular interplay.

In summary, KiMONo is a versatile method to derive fully integrated and holistic multi-level networks capturing the data-supported interplay between omics levels. Comprehensive benchmarks demonstrated that KiMONo is more sensitive to noise than to the reduction of samples. Further, application to two human disease settings showed that key nodes of the inferred multi-omics disease networks, also play key roles in disease pathophysiologies. Ultimately, the holistic networks inferred using KiMONo may serve as tools to easily uncover key regulatory features.

## References

Abelaira, Helena M., Gislaine Z. Réus, Morgana V. Neotti, and João Quevedo. 2014. “The Role of mTOR in Depression and Antidepressant Responses.” Life Sciences 101 (1-2): 10–14.

Anderson, Kevin M., Meghan A. Collins, Ru Kong, Kacey Fang, Jingwei Li, Tong He, Adam M. Chekroud, B. T. Thomas Yeo, and Avram J. Holmes. 2020. “Convergent Molecular, Cellular, and Neural Signatures of Major Depressive Disorder.” bioRxiv. https://doi.org/10.1101/2020.02.10.942227.

Argelaguet, Ricard, Britta Velten, Damien Arnol, Sascha Dietrich, Thorsten Zenz, John C. Marioni, Wolfgang Huber, Florian Buettner, and Oliver Stegle. n.d. “Multi-Omics Factor Analysis - a Framework for Unsupervised Integration of Multi-Omic Data Sets.” https://doi.org/10.1101/217554.

Arloth, Janine, Daniel M. Bader, Simone Röh, and Andre Altmann. 2015. “Re-Annotator: Annotation Pipeline for Microarray Probe Sequences.” PloS One 10 (10): e0139516.

Arloth, Janine, Ryan Bogdan, Peter Weber, Goar Frishman, Andreas Menke, Klaus V. Wagner, Georgia Balsevich, et al. 2015. “Genetic Differences in the Immediate Transcriptome Response to Stress Predict Risk-Related Brain Function and Psychiatric Disorders.” Neuron 86 (5): 1189–1202.

Breen, Michael S., Aliza P. Wingo, Nastassja Koen, Kirsten A. Donald, Mark Nicol, Heather J. Zar, Kerry J. Ressler, Joseph D. Buxbaum, and Dan J. Stein. 2018. “Gene Expression in Cord Blood Links Genetic Risk for Neurodevelopmental Disorders with Maternal Psychological Distress and Adverse Childhood Outcomes.” Brain, Behavior, and Immunity 73 (October): 320–30.

Bridge, Julia A., Janos Sumegi, Mihaela Druta, Marilyn M. Bui, Evita Henderson-Jackson, Konstantinos Linos, Michael Baker, Christine M. Walko, Sherri Millis, and Andrew S. Brohl. 2019. “Clinical, Pathological, and Genomic Features of EWSR1-PATZ1 Fusion Sarcoma.” Modern Pathology: An Official Journal of the United States and Canadian Academy of Pathology, Inc 32 (11): 1593–1604.

Catzavelos, Charles, Nandita Bhattacharya, Yee C. Ung, James A. Wilson, Luba Roncari, Charanjit Sandhu, Patricia Shaw, et al. 1997. “Decreased Levels of the Cell-Cycle Inhibitor p27Kip1 Protein: Prognostic Implications in Primary Breast Cancer.” Nature Medicine. https://doi.org/10.1038/nm0297-227.

Ceder, Rebecca, Ylva Haig, Marina Merne, Annette Hansson, Xi Zheng, Karin Roberg, Matthias Nees, et al. 2012. “Differentiation-Promoting Culture of Competent and Noncompetent Keratinocytes Identifies Biomarkers for Head and Neck Cancer.” The American Journal of Pathology 180 (2): 457–72.

Duman, Ronald S., George K. Aghajanian, Gerard Sanacora, and John H. Krystal. 2016. “Synaptic Plasticity and Depression: New Insights from Stress and Rapid-Acting Antidepressants.” Nature Medicine 22 (3): 238–49.

Engle, E. C. 2010. “Human Genetic Disorders of Axon Guidance.” Cold Spring Harbor Perspectives in Biology. https://doi.org/10.1101/cshperspect.a001784.

Felix Krueger, Simon R. Andrews. 2011. “Bismark: A Flexible Aligner and Methylation Caller for Bisulfite-Seq Applications.” Bioinformatics 27 (11): 1571.

Gassen, Nils C., and Theo Rein. 2019. “Is There a Role of Autophagy in Depression and Antidepressant Action?” Frontiers in Psychiatry. https://doi.org/10.3389/fpsyt.2019.00337.

Guadagno, Elia, Michela Vitiello, Paola Francesca, Gaetano Calì, Federica Caponnetto, Daniela Cesselli, Simona Camorani, et al. 2017. “PATZ1 Is a New Prognostic Marker of Glioblastoma Associated with the Stem-like Phenotype and Enriched in the Proneural Subtype.” Oncotarget 8 (35): 59282–300.

Hanif, Farina, Kanza Muzaffar, Kahkashan Perveen, Saima M. Malhi, and Shabana U. Simjee. 2017. “Glioblastoma Multiforme: A Review of Its Epidemiology and Pathogenesis through Clinical Presentation and Treatment.” Asian Pacific Journal of Cancer Prevention: APJCP 18 (1): 3–9.

Hapak, Sophie M., Carla V. Rothlin, and Sourav Ghosh. 2019. “aPKC in Neuronal Differentiation, Maturation and Function.” Neuronal Signaling. https://doi.org/10.1042/ns20190019.

Hasin, Yehudit, Marcus Seldin, and Aldons Lusis. 2017. “Multi-Omics Approaches to Disease.” Genome Biology. https://doi.org/10.1186/s13059-017-1215-1.

Heinig, Matthias, Michiel E. Adriaens, Sebastian Schafer, Hanneke W. M. van Deutekom, Elisabeth M. Lodder, James S. Ware, Valentin Schneider, et al. 2017. “Natural Genetic Variation of the Cardiac Transcriptome in Non-Diseased Donors and Patients with Dilated Cardiomyopathy.” Genome Biology 18 (1): 170.

Howard, David M., Mark J. Adams, Toni-Kim Clarke, Jonathan D. Hafferty, Jude Gibson, Masoud Shirali, Jonathan R. I. Coleman, et al. 2019. “Genome-Wide Meta-Analysis of Depression Identifies 102 Independent Variants and Highlights the Importance of the Prefrontal Brain Regions.” Nature Neuroscience 22 (3): 343–52.

Huang, Sijia, Kumardeep Chaudhary, and Lana X. Garmire. 2017. “More Is Better: Recent Progress in Multi-Omics Data Integration Methods.” Frontiers in Genetics 8 (June): 84.

Hua, Yunfeng, Andrew Woehler, Martin Kahms, Volker Haucke, Erwin Neher, and Jürgen Klingauf. 2013. “Blocking Endocytosis Enhances Short-Term Synaptic Depression under Conditions of Normal Availability of Vesicles.” Neuron. https://doi.org/10.1016/j.neuron.2013.08.010.

Ignácio, Zuleide M., Gislaine Z. Réus, Camila O. Arent, Helena M. Abelaira, Meagan R. Pitcher, and João Quevedo. 2016. “New Perspectives on the Involvement of mTOR in Depression as Well as in the Action of Antidepressant Drugs.” British Journal of Clinical Pharmacology 82 (5): 1280–90.

Kandaswamy, Radhika, Andrew McQuillin, David Curtis, and Hugh Gurling. 2012. “Tests of Linkage and Allelic Association between Markers in the 1p36 PRKCZ (protein Kinase C Zeta) Gene Region and Bipolar Affective Disorder.” American Journal of Medical Genetics. Part B, Neuropsychiatric Genetics: The Official Publication of the International Society of Psychiatric Genetics 159B (2): 201–9.

Kapur, S., A. G. Phillips, and T. R. Insel. 2012. “Why Has It Taken so Long for Biological Psychiatry to Develop Clinical Tests and What to Do about It?” Molecular Psychiatry. https://doi.org/10.1038/mp.2012.105.

Krumsiek, Jan, Karsten Suhre, Thomas Illig, Jerzy Adamski, and Fabian J. Theis. 2011. “Gaussian Graphical Modeling Reconstructs Pathway Reactions from High-Throughput Metabolomics Data.” BMC Systems Biology 5 (January): 21.

Lawrence, Michael, Wolfgang Huber, Hervé Pagès, Patrick Aboyoun, Marc Carlson, Robert Gentleman, Martin T. Morgan, and Vincent J. Carey. 2013. “Software for Computing and Annotating Genomic Ranges.” PLoS Computational Biology 9 (8): e1003118.

Lee, Jason D., and Trevor J. Hastie. 2015. “Learning the Structure of Mixed Graphical Models.” Journal of Computational and Graphical Statistics: A Joint Publication of American Statistical Association, Institute of Mathematical Statistics, Interface Foundation of North America 24 (1): 230–53.

Liu, Jing, Jinling Xu, Huining Li, Cuiyun Sun, Lin Yu, Yanyan Li, Cuijuan Shi, et al. 2015. “miR-146b-5p Functions as a Tumor Suppressor by Targeting TRAF6 and Predicts the Prognosis of Human Gliomas.” Oncotarget 6 (30): 29129–42.

McCord, Matthew, Yoh-Suke Mukouyama, Mark R. Gilbert, and Sadhana Jackson. 2017. “Targeting WNT Signaling for Multifaceted Glioblastoma Therapy.” Frontiers in Cellular Neuroscience. https://doi.org/10.3389/fncel.2017.00318.

Ogris, Christoph, Thomas Helleday, and Erik L. L. Sonnhammer. 2016. “PathwAX: A Web Server for Network Crosstalk Based Pathway Annotation.” Nucleic Acids Research 44 (W1): W105–9.

Ohta, Tomohiko, and Mamoru Fukuda. 2004. “Ubiquitin and Breast Cancer.” Oncogene. https://doi.org/10.1038/sj.onc.1207371.

Oughtred, Rose, Chris Stark, Bobby-Joe Breitkreutz, Jennifer Rust, Lorrie Boucher, Christie Chang, Nadine Kolas, et al. 2019. “The BioGRID Interaction Database: 2019 Update.” Nucleic Acids Research 47 (D1): D529–41.

Pinu, Farhana R., David J. Beale, Amy M. Paten, Konstantinos Kouremenos, Sanjay Swarup, Horst J. Schirra, and David Wishart. 2019. “Systems Biology and Multi-Omics Integration: Viewpoints from the Metabolomics Research Community.” Metabolites 9 (4). https://doi.org/10.3390/metabo9040076.

Relja, Borna, Katharina Mörs, and Ingo Marzi. 2018. “Danger Signals in Trauma.” European Journal of Trauma and Emergency Surgery: Official Publication of the European Trauma Society 44 (3): 301.

Roberson-Nay, Roxann, Aaron R. Wolen, Dana M. Lapato, Eva E. Lancaster, Bradley T. Webb, Bradley Verhulst, John M. Hettema, and Timothy P. York. n.d. “Twin Study of Early-Onset Major Depression Finds DNA Methylation Enrichment for Neurodevelopmental Genes.” https://doi.org/10.1101/422345.

Ronen, Jonathan, Sikander Hayat, and Altuna Akalin. 2019. “Evaluation of Colorectal Cancer Subtypes and Cell Lines Using Deep Learning.” Life Science Alliance 2 (6). https://doi.org/10.26508/lsa.201900517.

Sass, Steffen, Adriana Pitea, Kristian Unger, Julia Hess, Nikola S. Mueller, and Fabian J. Theis. 2015. “MicroRNA-Target Network Inference and Local Network Enrichment Analysis Identify Two microRNA Clusters with Distinct Functions in Head and Neck Squamous Cell Carcinoma.” International Journal of Molecular Sciences 16 (12): 30204–22.

Sinkala, Musalula, Nicola Mulder, and Darren Martin. 2020. “Machine Learning and Network Analyses Reveal Disease Subtypes of Pancreatic Cancer and Their Molecular Characteristics.” Scientific Reports 10 (1): 1212.

Terracciano, Antonio, Toshiko Tanaka, Angelina R. Sutin, Serena Sanna, Barbara Deiana, Sandra Lai, Manuela Uda, et al. 2010. “Genome-Wide Association Scan of Trait Depression.” Biological Psychiatry. https://doi.org/10.1016/j.biopsych.2010.06.030.

Varinthra, Peeraporn, and Ingrid Y. Liu. 2019. “Molecular Basis for the Association between Depression and Circadian Rhythm.” Ci Ji Yi Xue Za Zhi = Tzu-Chi Medical Journal 31 (2): 67–72.

Wang, Chun-Lei, Fu-Lei Tang, Yun Peng, Cheng-Yong Shen, Lin Mei, and Wen-Cheng Xiong. 2012. “VPS35 Regulates Developing Mouse Hippocampal Neuronal Morphogenesis by Promoting Retrograde Trafficking of BACE1.” Biology Open 1 (12): 1248–57.

Wang, X., N. Gulbahce, and H. Yu. 2011. “Network-Based Methods for Human Disease Gene Prediction.” Briefings in Functional Genomics. https://doi.org/10.1093/bfgp/elr024.

Weinstein, John N., The Cancer Genome Atlas Research Network, Eric A. Collisson, Gordon B. Mills, Kenna R. Mills Shaw, Brad A. Ozenberger, Kyle Ellrott, Ilya Shmulevich, Chris Sander, and Joshua M. Stuart. 2013. “The Cancer Genome Atlas Pan-Cancer Analysis Project.” Nature Genetics. https://doi.org/10.1038/ng.2764.

Yeo, Winnie, Paul K. S. Chan, Pun Hui, Wing M. Ho, Kwok C. Lam, Wing H. Kwan, Sheng Zhong, and Philip J. Johnson. 2003. “Hepatitis B Virus Reactivation in Breast Cancer Patients Receiving Cytotoxic Chemotherapy: A Prospective Study.” Journal of Medical Virology 70 (4): 553–61.

Zannas, Anthony S., Janine Arloth, Tania Carrillo-Roa, Stella Iurato, Simone Röh, Kerry J. Ressler, Charles B. Nemeroff, et al. 2015. “Lifetime Stress Accelerates Epigenetic Aging in an Urban, African American Cohort: Relevance of Glucocorticoid Signaling.” Genome Biology 16 (1): 1–12.

Zhernakova, Daria V., Patrick Deelen, Martijn Vermaat, Maarten van Iterson, Michiel van Galen, Wibowo Arindrarto, Peter van’t Hof, et al. 2017. “Identification of Context-Dependent Expression Quantitative Trait Loci in Whole Blood.” Nature Genetics 49 (1): 139–45.

Zierer, Jonas, Tess Pallister, Pei-Chien Tsai, Jan Krumsiek, Jordana T. Bell, Gordan Lauc, Tim D. Spector, Cristina Menni, and Gabi Kastenmüller. 2016. “Exploring the Molecular Basis of Age-Related Disease Comorbidities Using a Multi-Omics Graphical Model.” Scientific Reports 6 (November): 37646.

