## Supplementary material for "Knowledge guided multi-level network inference": Sup Fig

### Supplemental Figures & Tables

| Gene symbol | #Publication hits | Gene symbol | #Publication hits |
|---|---|---|---|
| BRCA1 | 271 000 | HNRNPA1 | 5 370 |
| CDK2 | 74 300 | HSP90AA1 | 4 410 |
| DLG4 | 1 540 | MYC | 1 030 000 |
| EP300 | 11 100 | SKP1 | 14 600 |
| FBXW11 | 844 | UBC | 45 600 |
| GRB2 | 47 300 | UBE2I | 1 630 |
| HDAC1 | 41 600 | AKT1 | 52 400 |

*Sup. Table 1: KiMONo identified 14 genes which are identified as important across all 11 panCancer networks. The publication hits define the approximate amount of publication including the gene. These hits were derived using google scholar searches.*

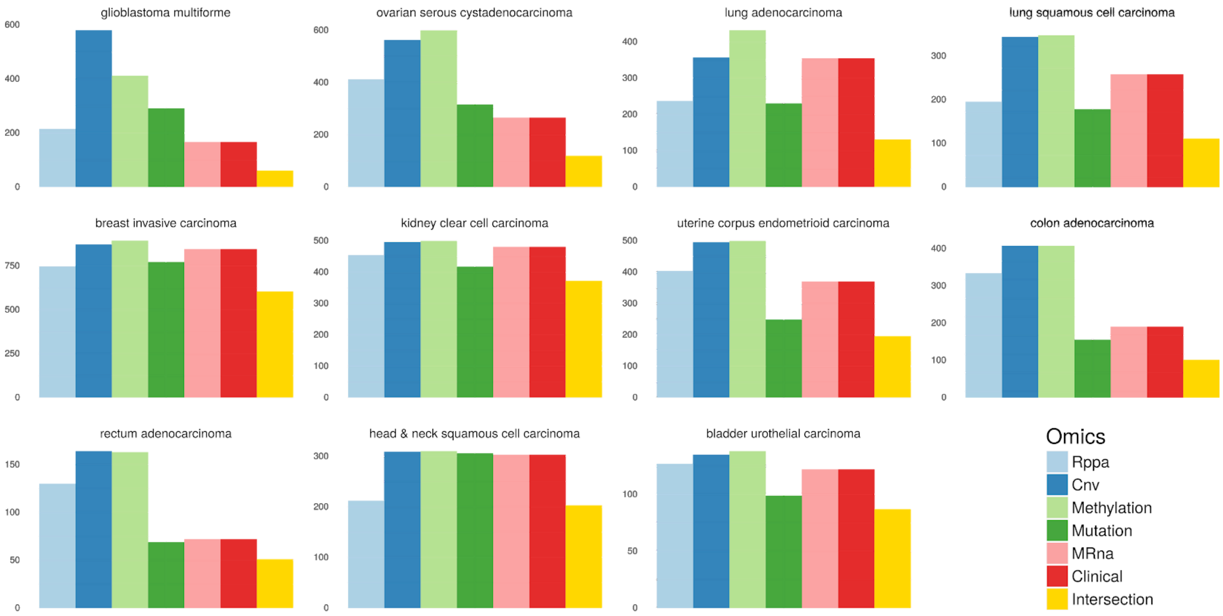


***Sup. Figure 1:*** *Overview of pan cancer data for 11 different cancer types and 6 different data types. Proteomics / Rppa (blue), Copy number variations / CNV (dark blue), Methylation (limegreen), Mutations (dark green), mRNA (pink) and Clinical information (red). The intersection (yellow) denotes the amount of matched samples. In our analysis we only used the samples which were analysed across all levels.*

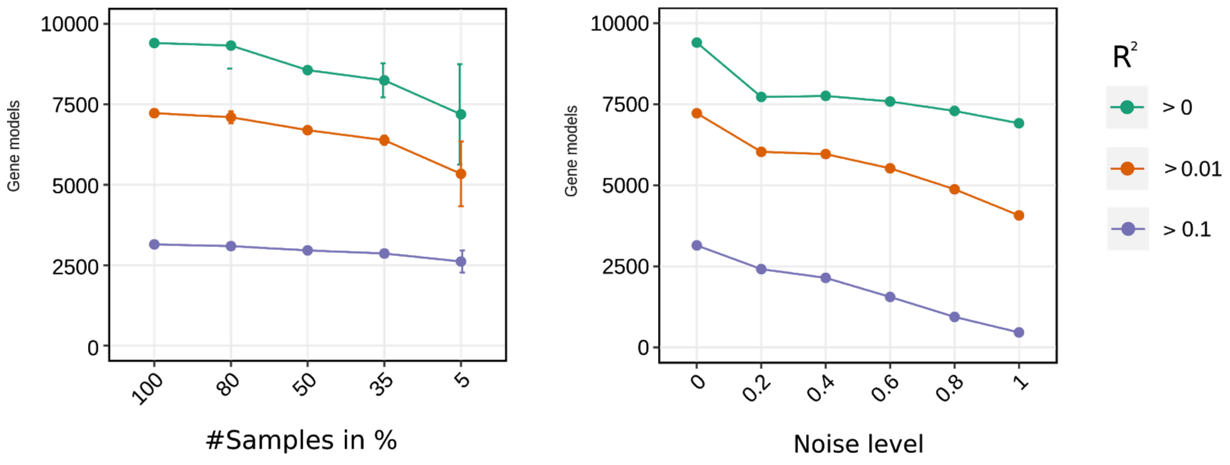


***Sup. Figure 2:*** *Overview of network coverage for both benchmarks.*

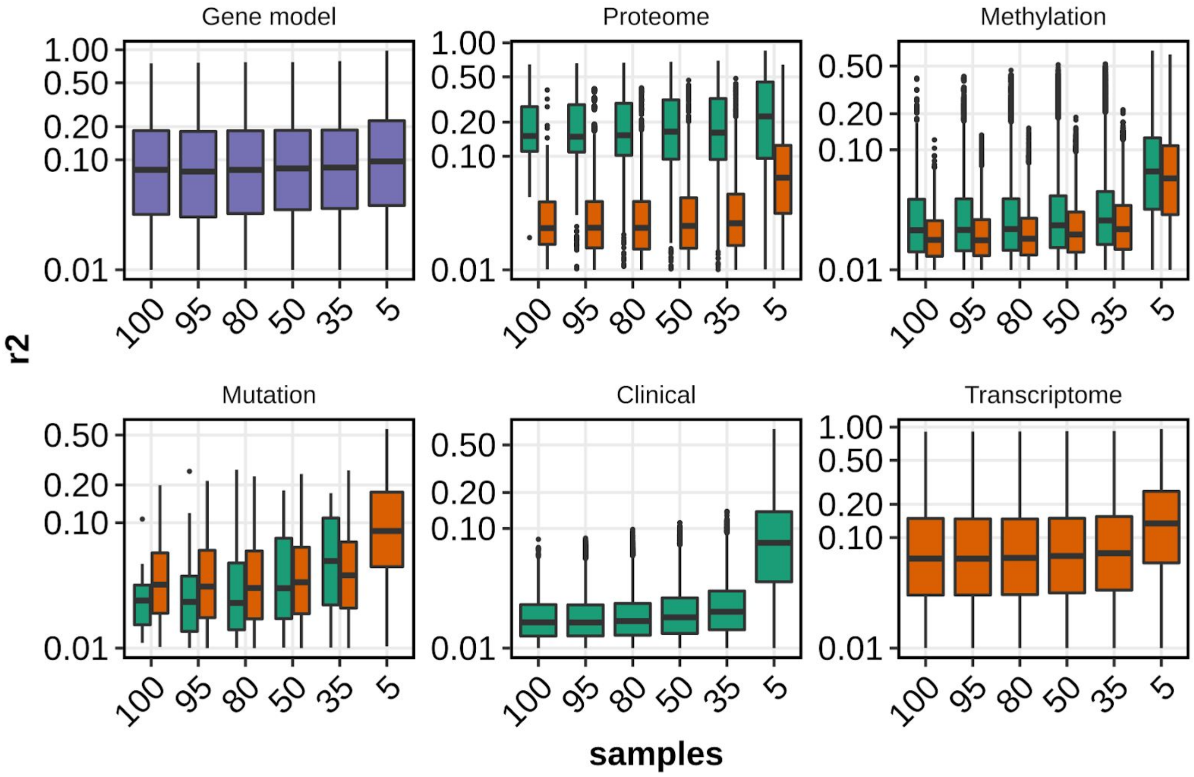


***Sup. Figure 3:*** *Results of benchmarking small sample sizes and different noise levels. Here, we used all inferred models which explain at least 1% of the variance in the data.*

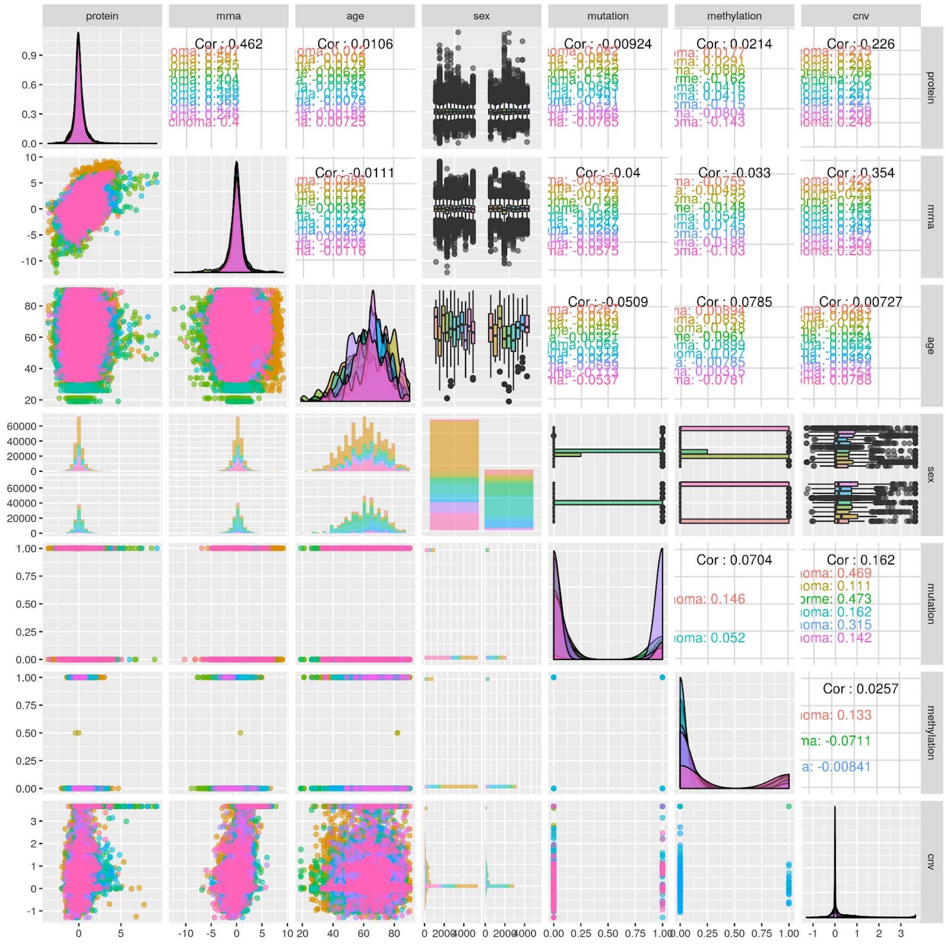


***Sup. Figure 4:*** *Overview of the available TCGA raw omic data for all 10 cancer types. Rows and columns denote the omic levels. For the clinical data level each feature is visualized separately since it consisted of binarized and continuous data.*

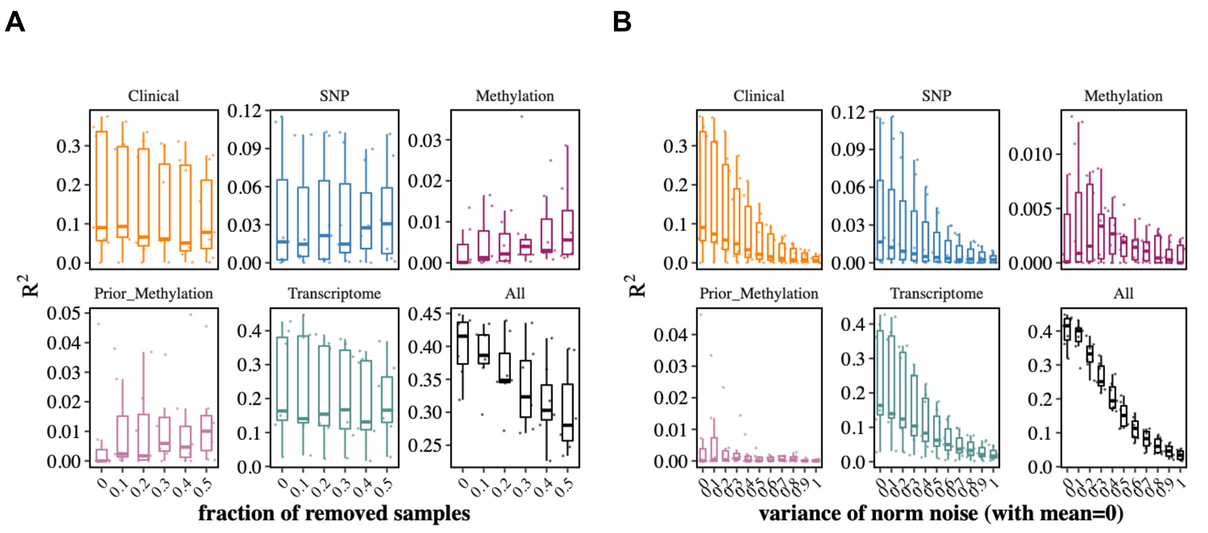


***Sup Figure 5:*** *Robustness benchmark for A) different sample sizes and B) noise levels on MDD data. The boxplots show the performance* $R^2$ *of inferred gene models. Panels describing the performance of stand-alone first-order links are displayed first (Clinical, SNPs and Methylation), followed by second-order links (Prior Methylation and Transcriptome). The last panel shows the performance of inferred gene models using all available information layers. A) Data sets with different sample sizes were generated using 10% - 50% of the 107 MDD samples. B) Different test data sets were simulated by adding Gaussian noise with increasing variance. Here, the noise level reflects the* $\sigma$ *for ten intensities.*
